# Long-term effects of psilocybin on dynamic and effectivity connectivity of fronto-striatal-thalamic circuits

**DOI:** 10.1101/2024.11.06.622302

**Authors:** Lorenzo Pasquini, Jakub Vohryzek, Anira Escrichs, Yonatan Sanz Perl, Adrian Ponce-Alvarez, Sebastian Idesis, Manesh Girn, Leor Roseman, Jennifer M. Mitchell, Adam Gazzaley, Morten Kringelbach, David J. Nutt, Taylor Lyons, Robin L. Carhart-Harris, Gustavo Deco

## Abstract

Psilocybin has been shown to induce fast and sustained improvements in mental well-being across various populations, yet its long-term mechanisms of action are not fully understood. Initial evidence suggests that longitudinal functional and structural brain changes implicate fronto-striatal-thalamic (FST) circuitry, a broad system involved in goal-directed behavior and motivational states. Here, we apply empirical methods and computational modeling to resting-state fMRI data from a within-subject longitudinal psilocybin trial in psychedelic-naïve healthy volunteers. We first show increases in FST dynamic activity four weeks after a full dose of psilocybin. We then proceed to mechanistically account for these increased dynamics, by showing that reduced structural constraints underlie increased FST dynamic activity post psilocybin. Further, we show that these reduced structural constraints come along with increased bottom-up and reduced top-down modulation of FST circuits. While cortical reductions in top-down modulation are linked to regional 5-HT2A receptor availability, increased information outflow via subcortical and limbic regions relate to local D2 receptor availability. Together, these findings show that increased FST flexibility weeks after psilocybin administration is linked to serotonergic-mediated decreases in top-down information flow and dopaminergic-mediated increases in bottom-up information flow. This long-term functional re-organization of FST circuits may represent a common mechanism underling the potential clinical efficacy of psilocybin across various neuropsychiatric disorders including substance abuse, major depression, and anorexia.

**Significance Statement:** Fronto-striatal-thalamic systems, which underlie motivation and reward, go through profound functional and structural changes following psilocybin administration. We leveraged longitudinal fMRI data from a within-subject psilocybin trial in psychedelic-naïve healthy participants to show that psilocybin increases fronto-striatal-thalamic dynamic activity as well as flexibility four weeks after dosing. Computational modeling revealed that this increased flexibility is mechanistically caused by reduced structural constraints on functional dynamics. Further long-term changes included increased bottom-up and reduced top-down information flow mediated by the serotonergic and dopaminergic systems. This long-term functional re-organization of fronto-striatal-thalamic circuits may reflect a common mechanism underlying clinical symptoms improvements across diagnostic groups, such as increased openness, improved well-being, and reductions in anhedonia, apathy, and substance craving.

## Introduction

Psilocybin is currently being explored as a potential treatment for a variety of highly debilitating neuropsychiatric disorders, including depression, anxiety, addiction, and eating disorders (1–7). Several studies have shown fast improvements in clinical measures and mental well-being, revealing enduring effects for up to six months after one-to-two moderate-to-high psilocybin doses (1–7). These remarkable findings compel the exploration of neural mechanism underlying such profound long-term changes across these various clinical populations, which range from alleviation of anhedonia and apathy (2–4), to reduced substance craving (5, 6), and increased openness (8).

The serotonin 5-HT2A receptor is widely regarded as the key site for the acute action of psychedelic drugs, including psilocybin (9). The 5-HT2A receptor is highly expressed among higher-level cognitive, heteromodal brain areas overlapping with brain networks supporting attentional, self-referential, and emotional processes (10). Under acute effects, psilocybin has been shown to induce reduced integration within heteromodal and higher-cognitive brain networks (11, 12). These circumscribed reductions in network cohesion go along with higher global between-network integration, with such functional changes correlating with the intensity of drug effects (11, 12). Although neuroimaging has played a crucial role in elucidating psilocybin’s acute effects on the brain, only a handful of studies have attempted to explore the long-term changes induced by psilocybin and related psychedelics on brain function of healthy and clinical populations (1, 13–15). These studies consistently report increases in global brain network integration taking place days to weeks after drug administration, with changes in connectivity correlating with longitudinal increases in well-being and with the intensity of the altered state of consciousness under the acute drug effects.

Common findings across studies include marked functional changes among a set of midline frontal brain areas anchored by the prefrontal and anterior cingulate cortices (1, 13, 14, 16). Recently, these regions have also been shown to undergo white matter changes, with a previous analysis on the same sample analyzed here revealing decreased axial diffusivity in prefrontal-subcortical tracts following a dose of 25 mg psilocybin (17). These tracts are core components of fronto-striatal-thalamic (FST) circuits, which mostly originate in the prefrontal cortex and project to the striatum, globus pallidus, substantia nigra, and finally to the thalamus, where feedback loops project back to the prefrontal cortices (18). FST circuits receive inputs from major modulatory systems, including the serotoninergic and dopaminergic systems, overlap with the mesolimbic reward pathway (18, 19), and have been implicated in a variety of processes ranging from executive functions, affective and motivational states, pain processing, social cognition, future planning, and reward prediction (18, 20). Due to their role in goal-directed behavior and to their input from major neuromodulatory systems, FST circuits represent a potential target for the long-term, cross-diagnostic motivational and affective changes taking place after psilocybin administration (21–23).

Generative computational approaches provide a powerful biophysical framework to model both emerging functional brain dynamics and directed information flow between brain regions as a function of underlying anatomical constraints (24–26). These models leverage mathematical concepts used to formalize dynamic systems and oscillators and, through the modulation of few parameters, simulate activity time series that reproduce the statistical properties of empirical brain activity (24–26). Whole-brain models have been successfully applied to resting-state fMRI data to describe the emergence of brain states across distinct clinical conditions (e.g., stroke or Alzheimer’s disease)(27, 28) and among distinct levels of consciousness (e.g., minimally conscious states, sleep stages, awake, acute psychedelics)(29–32).

Recently, these generative models have also been extended to elucidate the mechanisms underlying hierarchical causal interactions between brain regions across various conditions, including movie watching versus rest (33), or various states of consciousness (awake, deep sleep, anesthesia)(34). This approach borrows concepts from thermodynamics and estimates the signal propagation hierarchy between brain regions by assessing the temporal asymmetry, or reversibility, of brain signals (33–35). The degree of asymmetry is assessed using a time-shifted measure of the correlations between the forward and the reversed time series. The difference between these time-shifted correlations quantifies the asymmetry in the interactions between pairs of brain areas representing the extent to which one area influences another. Consequently, it unveils the causal mechanisms responsible for generating specific dynamic changes in brain states.

Here, we capitalize on these approaches to mechanistically account for long-term changes in FST dynamic activity and to elucidate changes in causal interactions between FST regions taking place four weeks after 25 mg of psilocybin. To reach these aims, we apply empirical methods as well as computational models (24–26) to resting-state fMRI data from a longitudinal, within-subject psilocybin trial in healthy, psychedelic-naïve volunteers (17).

## Results

### Increased dynamicity of FST activity following 25 mg psilocybin

We assessed the effects of psilocybin on FST functional dynamics by capitalizing on resting-state fMRI data acquired in psychedelic-naïve healthy adults (***Table S1***) at three times: at baseline (B1), four weeks after a control dose of 1 mg psilocybin (B2), and four weeks after a full active dose of 25 mg psilocybin (key endpoint KE, **Figure 1A, *Figure S1***). Blood-oxygen-level-dependent (BOLD) activity time-series were extracted from 28 FST regions-of-interest (**Figure 1B-C**) and used to measure dynamic functional homogeneity (DFH, **Figure 1D**) for each individual at each scan. DFH was computed by assessing the similarity of activity over different temporal periods and provides an index of FST activity homogeneity over time, as described in ***Materials and Methods***. We then investigated whether DFH, where lower values indicate increased activity heterogeneity and higher levels of FST functional dynamics, differed across the three scanning time points (**Figure 1E**). Mean FST DFH did not significantly differ when comparing the baseline to the post-1 mg psilocybin scan (*t(24)* = 0.26, *p* = 0.79). Mean FST DFH acquired post-25 mg psilocybin was significantly lower than DFH post-1 mg psilocybin (*t(24)* = 2.87, *p* < 0.01) and trending lower than DFH at baseline (*t(24)* = 1.92, *p* < 0.1). Crucially, reductions in DFH post-25 mg were specific to the FST and not observed at the whole-brain level (***Figure S2***). Reductions in FST DFH between the second and third scan correlated with mental well-being increases assessed in the same time interval i.e., one month post-25 mg psilocybin (**Figure 1F**, Rho*(22)* = -0.36, *p* < 0.1). Correlations were corrected for baseline mental well-being and differences in mean head frame-wise displacement between both scans. These findings suggest longitudinal increases in FST dynamic activity following psilocybin administration, with such increases contributing to mental well-being improvements after dosing. Finally, we used computational modelling to investigate whether psilocybin mechanistically alters the coupling between structural connectivity and FST dynamic activity. We used a partial-brain model, a variant of the regular whole-brain Hopf model (24–26) (***Figure S3A***), to generate simulated BOLD activity time series reproducing (***Figure S3B***) the statistical properties of empirical brain dynamics (***Figure S3C-D***). Few parameters are needed to fit the model, with the global coupling parameter *G* reflecting the constraining effect of structural connectivity on emerging functional dynamics. We found a lower optimal *G* value for FST dynamics post-25 mg psilocybin when compared to post-1 mg psilocybin (***Figure S3E***) with significantly lower fitting metrics (***Figure S3F***, *z(29)* = 6.60, *p* < 0.0005). Low values of *G* reflect instances where FST dynamic activity is weakly constrained by the underlying structural connections (36, 37), hence reflecting uncoupling of FST dynamics from the underlying structural constraints post-25 mg psilocybin.

**Figure 1.**
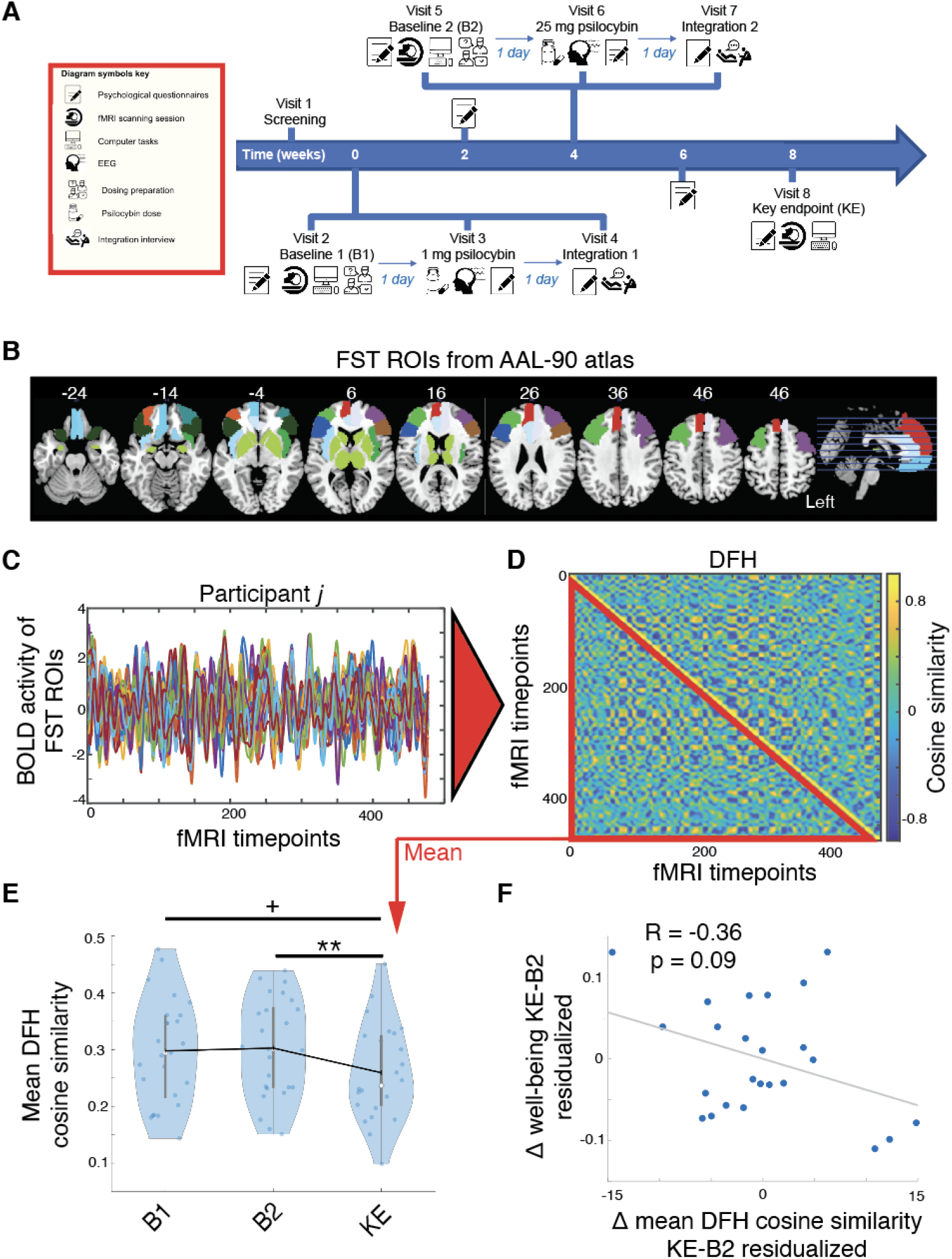
Decreased FST dynamic functional homogeneity following 25 mg psilocybin. **(A)** Within-subject study design. 25 psychedelic-naïve healthy volunteers were dosed with 1 mg psilocybin one day after receiving an fMRI and surveys at baseline (B1). Volunteers received a second fMRI and surveys (B2) four weeks after this first dosing session, followed by 25 mg psilocybin the day after. A last fMRI assessment and key endpoint surveys (KE) were completed four weeks after being dosed with 25 mg psilocybin. **(B)** FST regions-of-interest (ROIs) from the AAL-90 atlas. **(C)** Time-series of resting-state BOLD activity extracted from the FST ROIs shown for an individual participant. **(D)** FST activity time-series were used to generate a time-resolved cosine similarity matrix of FST activity for each participant, reflecting FST dynamic functional homogeneity (DFH). Lower values of DFH reflect higher heterogeneity of FST activity across a scanning session. The lower triangular matrix was then averaged to derive an individual estimate of FST DFH for each participant. **(E)** Mean cosine similarity of FST DFH was significantly reduced at KE four weeks after dosing with 25 mg psilocybin, when compared to B2 assessed four weeks after a control dosing with 1 mg psilocybin and when compared to B1 assessed before dosing. **(F)** Reduction in FST DFH at KE compared to B2 correlated with increases in mental well-being, assessed through the Warwick-Edinburgh Mental Well-being Scale. The influence of well-being at baseline and change in mean head frame-wise displacement was regressed out before performing the correlation. Panel A adapted with permission from (17). +*p* < 0.1; \*\**p* < 0.01. FST = fronto-striatal-thalamic.

### Psilocybin alters effective connectivity between FST nodes

While in the previous section we used empirical and computational methods to show increased FST dynamic activity following a high dose of psilocybin, we next aimed to assess whether these dynamic changes are accompanied by mechanistic change in effective connectivity. To do so we leveraged a whole-brain model that capitalizes on the underlying structural connectivity (**Figure 2A**) and the inherent time-dependent asymmetry of brain signals to estimate the directionality of information flow between brain regions. Anatomical constraining parameters are used to derive the causal interaction between brain regions by maximizing the fit between the empirical and the simulated intrinsic functional connectivity and time-shifted forward and reversed covariance matrices (see *Materials and Methods* for more detail; **Figure 2B**). This approach yielded effective connectivity matrices estimated separately for the post-1mg and the post-25 mg psilocybin scans, as well as a subtraction matrix comparing both post-dosing sessions (**Figure 2C-E**, *p* < 0.05 FDR corrected). Averaging across rows provides an estimate of outflowing effective connectivity (EC-out), or total information flow from the source region to target regions, while averaging across columns provides an estimate of total inflowing effective connectivity (EC-in), or total information flow from source regions to the target region. Outflowing effective connectivity following 25 mg psilocybin was reduced among dorsal cortical regions spanning the dorsolateral prefrontal and medial frontal cortices, while it was increased in limbic and subcortical areas (**Figure 2F**). Inflowing effective connectivity following 25 mg psilocybin showed a similar pattern, with reductions among the dorsolateral prefrontal and medial frontal cortices, yet, it was increased among limbic and subcortical areas, particularly on the left hemisphere (**Figure 2G**). Nodal changes in outflowing effective connectivity following 25 mg psilocybin correlated with changes in inflowing effective connectivity (**Figure 2H**, Rho(*26*) = 0.80, *p* < 0.0005), reflecting long-term increases in bottom-up modulation along with reduced top-down contributions following 25 mg psilocybin.

**Figure 2.**
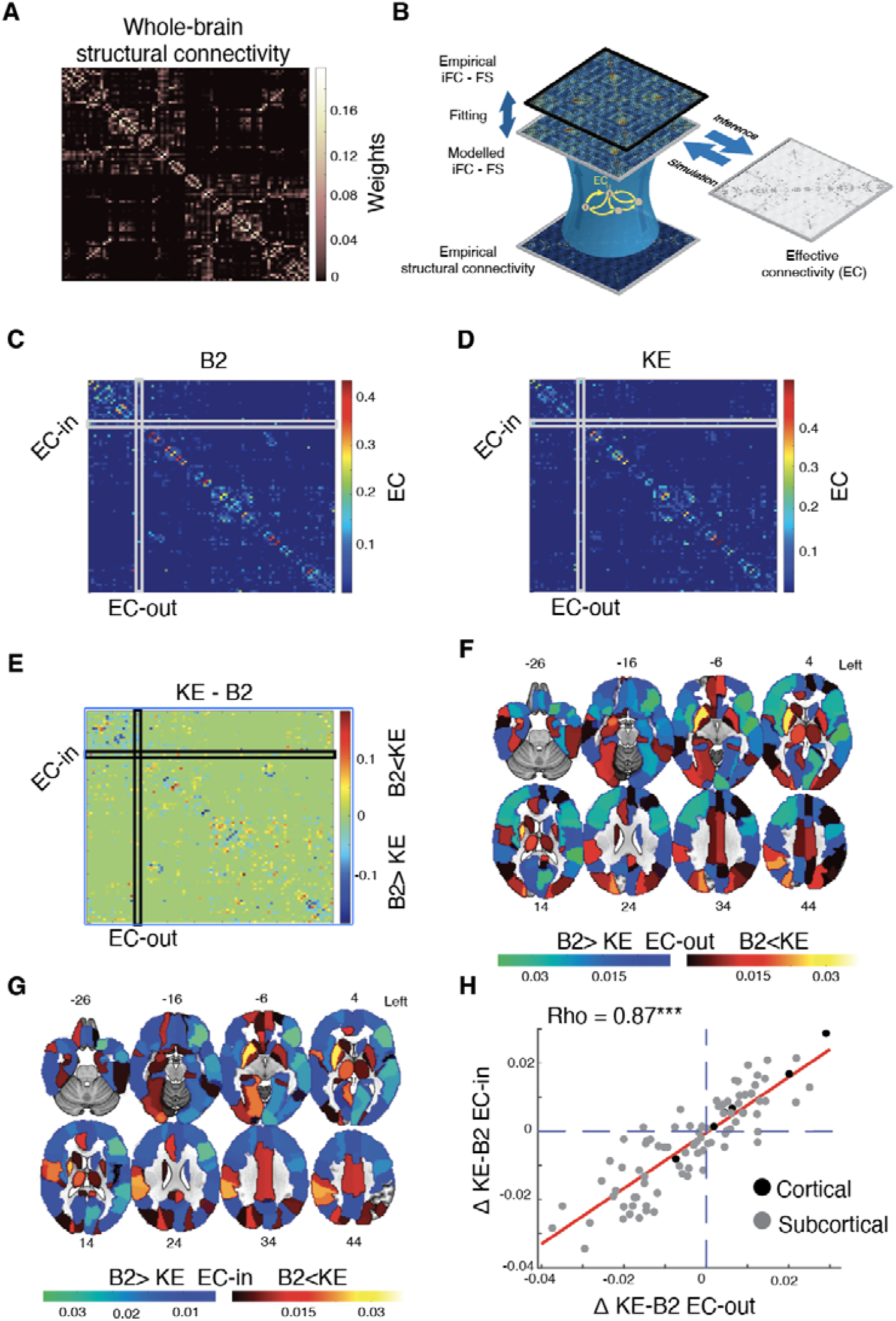
Whole-brain effective connectivity after 25 mg psilocybin. **(A)** Structural connectivity matrix derived from diffusion tensor imaging data of a normative healthy control sample. **(B)** We used a whole-brain model leveraging the underlying structural connections to derive a measure of effective connectivity (EC), reflecting the directional flow of information between pairs of brain regions. EC between pairs of brain regions is computed through an iterative process optimizing the distance between modelled and empirical difference of intrinsic functional connectivity (iFC) and time-shifted covariance matrices (FS). The model is run repeatedly with the updated effective connections until convergence. **(C)** EC between pairs of brain regions at B2, four weeks after 1 mg psilocybin. Averaging across rows provides an estimate of outflowing EC (EC-out), while averaging across columns provides an estimate of inflowing EC (EC-in). **(D)** EC between pairs of brain regions at KE, four weeks after 25 mg psilocybin. **(E)** Heatmap reflecting significant increases (warm colors) and decreases (cold colors) in EC between pairs of brain regions when comparing post-25 mg to post-1 mg psilocybin (*p* < 0.05 FDR corrected for multiple comparisons). Nodal increases (warm colors) and decreases (cold colors) in EC-out **(F)** and EC-in **(G)** when comparing post-25 mg to post-1 mg psilocybin. **(H)** Nodal changes in EC-out following 25 mg psilocybin positively correlate with nodal changes in EC-in after 25 mg psilocybin. Panel **B** adapted with permission from (30). \*\*\**p* < 0.0005.

### FST effective connectivity changes correlate with local 5-HT2A receptor and D2 transporter density

The acute effects of psilocybin are primarily mediated through its agonist action on the serotonergic 5-HT2A receptor (9). We wondered whether also long-term changes in causal interactions between brain regions related to the regional availability of 5-HT2A receptors. We derived regional FST maps of 5-HT2A receptor densities by leveraging publicly available maps from standardized radiotracer positron emission tomography (PET) studies (38). The FST 5-HT2A receptor density map (**Figure 3A**) was correlated with nodal maps of outflowing and inflowing effective connectivity change following 25 mg psilocybin administration (**Figure 3B-C**). These analyses were conducted separately for cortical and subcortical areas due to these structures showing marked differences in PET signal intensity values and due to opposing changes in inflowing and outflowing effective connectivity between subcortical and cortical areas. The distribution of the 5-HT2A receptor correlated negatively with both cortical inflowing (Rho(*18*) = -0.56, *p* < 0.05) and outflowing (Rho(*18*) = -0.47, *p* < 0.05) effective connectivity changes. No significant correlations were found between 5-HT2A receptor availability and subcortical inflowing (Rho(*6*) = 0.33, *p* = 0.42) or outflowing (Rho(*6*) = 0.21, *p* = 0.62) effective connectivity changes, with subcortical correlations being significantly different from cortical ones (**Figure 3B-C**). Crucially, whole-brain 5-HT2A receptor availability did not correlate with whole-brain cortical inflowing (Rho(*80*) = -0.11, *p* = 0.33) or outflowing (Rho(*80*) = -0.11, *p* = 0.33) effective connectivity changes, suggesting a specific association between FST effective connectivity changes following 25 mg psilocybin and 5-HT2A receptor availability.

**Figure 3.**
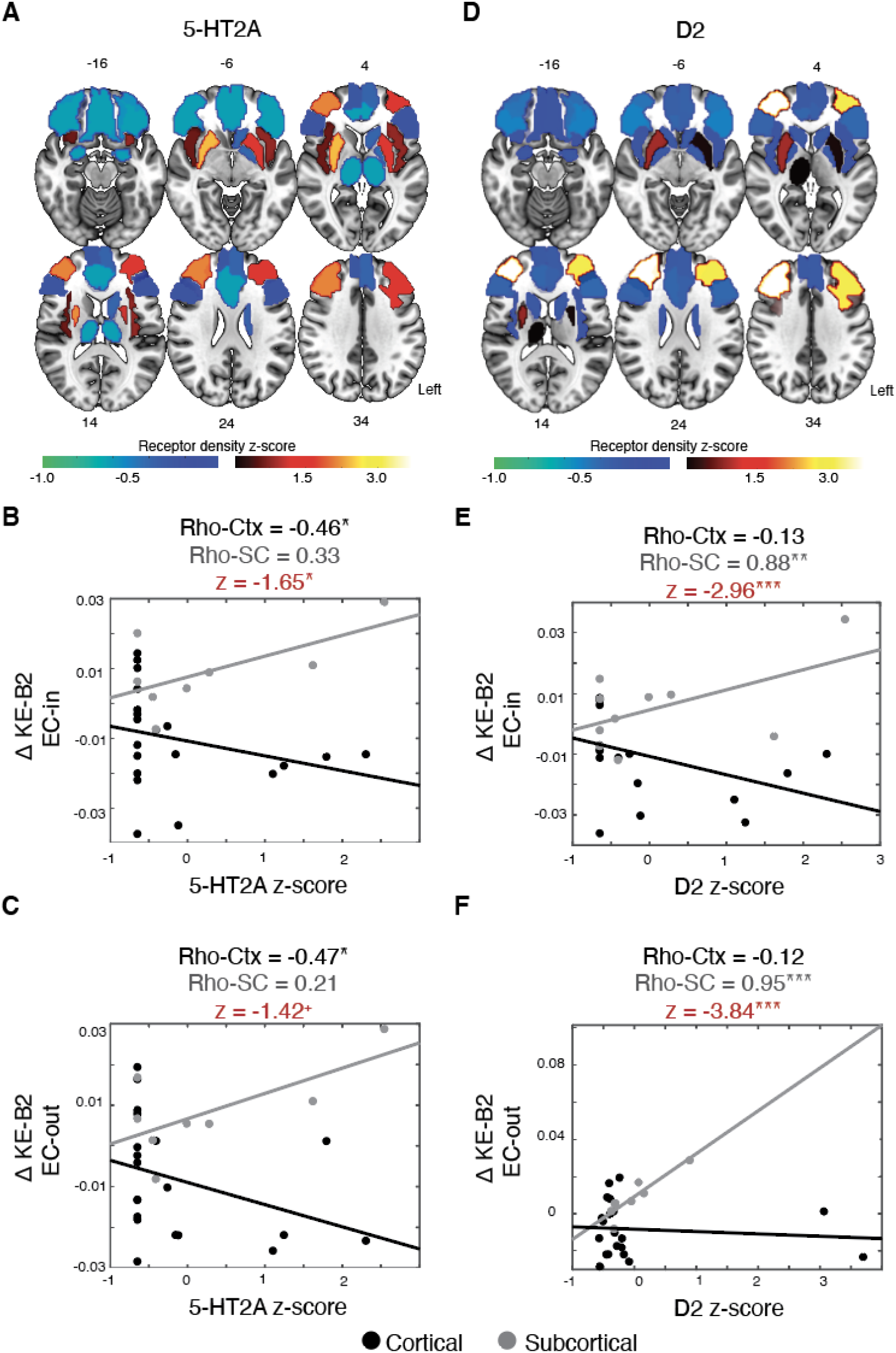
Effective connectivity changes correlate with regional densities of serotonin receptors and dopamine availability. Spatial distribution of 5-HT2A **(A)** serotonergic receptors within FST regions (z-scores), warmer colors reflect higher densities. Scatterplots reflecting the association between nodal 5-HT2A receptor densities and changes in EC-in **(B)** and EC-out **(C)** following 25 mg psilocybin, plotted separately for cortical (Ctx, black circles) and subcortical regions (SC; grey circles). **(D)** FST density map of the D2 dopamine receptor. Cortical and subcortical association between changes in EC-in **(E)** and EC-out **(F)** following 25 mg psilocybin and local D2 receptor density. Z-statistics (in red) test the difference in correlation strength across cortical and subcortical nodes. ^+^*p* < 0.1; \**p* < 0.05; \*\**p* < 0.01; \*\*\**p* < 0.005.

Given that our analyses failed to find a subcortical relationship between effective connectivity changes and 5-HT2A receptor density, we turned our attention to the dopaminergic system, an important modulator of FST circuits (18, 19). FST D2 receptor density (**Figure 3D**) correlated positively with subcortical inflowing (**Figure 3E**, Rho(*8*) = 0.88, *p* < 0.01) and outflowing (**Figure 3F**, Rho(*8*) = 0.95, *p* < 0.005) effective connectivity changes but not with cortical inflowing (Rho(*18*) = -0.13, *p* = 0.57) and outflowing (Rho(*18*) = -0.12, *p* = 0.61) effective connectivity, with cortical correlations being significantly different from subcortical ones (**Figure 3E-F)**. Overall, these findings suggest that diminished FST top-down modulation following psilocybin administration is linked to the 5-HT2A receptor, while long-term effects on bottom-up outflow may results from a more complex influence of both the serotonergic and dopaminergic systems on subcortical and cortical FST activity.

## Discussion

In this study, we investigated long-term changes in FST dynamic activity and effective connectivity before and one month after psilocybin administration. Longitudinal resting-state fMRI data from a within-subject psilocybin trial in psychedelic naïve healthy controls were examined using both empirical and computational approaches. We used DFH to index functional homogeneity of dynamic activity in FST circuits, assessed across the duration of an entire fMRI session. FST DFH did not differ when comparing baseline to one month after a control dose of 1 mg psilocybin, suggesting that FST dynamics display long-term, stable trait characteristics over a one month period. In contrast, FST DFH after 25 mg psilocybin dosing was reduced when compared to either baseline or one month after 1 mg psilocybin.

This finding suggests a long-term reconfiguration of FST circuitry following a full dose of psilocybin, indicating increases in dynamicity and heterogeneity of FST activity. Long-term increases in FST dynamic activity correlated with longitudinal improvements in mental well-being following 25 mg psilocybin. This association was robust against outliers and corrected for both baseline well-being levels and longitudinal changes in head movement while scanned. Overall, this finding suggests long-term flexibility increases in FST activity after a full dose of psilocybin, with these increases partially accounting for longitudinal improvements in mental well-being. Our findings are in line with previous reports showing longitudinal anterior cingulate and prefrontal cortex functional re-organization days to weeks after the administration of a psychedelic substance. For example, the psychedelic brew ayahuasca has been shown to increase local and global connectivity of the anterior cingulate one day after administration, with connectivity changes being associated with acute scores of altered somesthesia and affect (13). Echoing our findings, dynamic functional connectivity of the anterior cingulate cortex has been shown to increase in healthy controls two weeks following two moderate-to-high psilocybin dosing sessions (14). Further, resting-state fMRI data acquired in the context of two psilocybin-based depression trials revealed increased global functional integration (or decreased network modularity) across both studies.

Decreased network modularity correlated with improved symptom severity in both cases (1). High level fronto-parietal networks such as the default mode and executive networks were especially implicated. Finally, we partially replicated a previous finding on the same sample analyzed in this study, in that we observed a consistent relationship between improved mental health and decreased brain network modularity one month after 25 mg psilocybin. Interestingly, this effect correlated with decreases in bilateral axial diffusivity of prefrontal-subcortical tracts one month after 25 mg psilocybin as well as with reports of psychological insight (17). Taken together, these findings highlight the longitudinal impact that psilocybin has on brain functional organization, with FST circuitry playing an apparent role in mediating long-term clinical outcomes. Yet, which mechanisms may account for such increases in FST activity homogeneity? Our mechanistic modelling approach may provide an answer to this question. We used a partial-brain model, a variant of the regular whole-brain Hopf model (24–26), to mechanistically describe the influence of anatomical constrains on increased FST dynamic activity found four weeks after 25 mg psilocybin. We found that the global coupling parameter *G*, describing the coupling between emerging FST dynamics and the underlying structural connectome, is decreased following 25 mg psilocybin. This finding suggests that, in the long-term, psilocybin relaxes structure-function coupling among FST regions, therefore contributing to the emergence of more flexible FST states, possibly via functional neural plasticity effects on the frontal cortex (39–41).

Further computational models leveraging the concept of non-reversibility revealed long-term effects of psilocybin on effective connectivity between FST nodes. In this context, effective connectivity is referring to the directional flow of information between brain regions, as commonly derived through Granger causality or dynamic causal modeling (42, 43). Yet, it differs from these methods by accounting for the constraining effects of local structural weights on causal interactions emerging between node pairs (33). Decreases in both inflowing and outflowing effective connectivity following a high dose of psilocybin were found among high-level brain regions exerting top-down information-flow overlapping with the dorsolateral prefrontal, frontal, and insular cortices. Conversely, subcortical and limbic regions lower in the functional hierarchy and assumed to exert more bottom-up information flow – such as the thalamus, putamen, and anterior cingulate – showed marked increases in both inflowing and outflowing effective connectivity. While decreases in top-down effective connectivity were related to the cortical availability of the 5-HT2A receptor, increases in bottom-up information outflow related to subcortical D2 receptor availability.

How are these findings in keeping within major theoretical models of acute psychedelic drug action? One model posits that psilocybin and related psychedelics acutely disrupt cortico-striatal-thalamic circuits through 5-HT2A activation in the prefrontal cortex, which in turn disrupt GABAergic pallido-thalamic neurons resulting in thalamus disinhibition and inundation of the cortex with sensory information i.e., an impaired thalamic gating model (22). Evidence for this model has been provided by directional modelling of resting-state fMRI data, showing that lysergic acid diethylamide, a classic psychedelic akin to psilocybin, acutely decreases striatal input to the thalamus (44). As our present study did not look at the acute psychedelic state, it may therefore not be surprising that our findings of subcortical increases in inflowing and outflowing information are inconsistent with a thalamic gating model. This suggests that thalamic gating may not account for the long-term effects of psilocybin on FST circuits. Another model, known as ‘Relaxed Beliefs Under Psychedelics’ (REBUS), proposes that psychedelics disrupt the hierarchical organization of brain function by dysregulating statistical regularities in the spontaneous activity of higher-level cortical areas – causing less efficient top-down control of activity in functionally lower-level regions (45). In Bayesian terms, the REBUS model states that psychedelics cause a relaxation of precision weightings encoding prior beliefs, resulting in less confident assumptions. Relaxing the precision-weighting of priors encoded in higher-level, cortical areas, would in turn liberate activity in previously ‘controlled’ lower-level regions. Overall, our findings support this model, extending it beyond the originally proposed acute action to a more enduring sub-acute action (i.e., one month post-25 mg psilocybin). The present results also suggest that long-term increases in subcortical information outflow and inflow correlate with the regional expression of the dopamine transporter. This finding could be viewed as consistent with recent preclinical work showing that psychedelics reopen a critical period for learning social reward (46), as well as with the well-established principle that dopamine neurotransmission in midbrain-striatum-orbitofrontal cortex circuits is especially involved in reinforcement learning(19) and possibly, belief updating (47). Importantly, given the low addiction potential of psilocybin and related psychedelics (9), and that diminished striatal connectivity has been associated with vulnerability to drug addiction (48) and alcohol dependence (49), this dopaminergic involvement is unlikely to reflect a drug-dependency risk for psilocybin.

The following limitations need to be considered when interpreting the present findings. First, our mechanistic approach needs to be extended to include other psychedelic substances before it can be extrapolated to psychedelics more generally. Second, FST regions exhibit high functional-anatomic heterogeneity and future studies would benefit from the use of more fine-grained, individualized FST parcellations and structural connectivity estimates. Further work implementing gradual dosing-protocols with multiple neuroimaging assessments is needed to evaluate the dose- and time-dependency of FST functional re-organization across healthy and clinical populations. Nonetheless, our findings open up new avenues for the application of mechanistic approaches to elucidate the long-term impact of pharmacological interventions on the brain. Specifically, our data suggest that psilocybin increases FST dynamic activity and subcortical information outflow through cortical serotonergic and subcortical dopaminergic systems. This FST functional re-organization, in turn, may underly cross-diagnostic longitudinal behavioral changes, including increased openness and well-being, and decreased anhedonia, apathy, and substance craving after psychedelics (1–7). Further research is necessary to determine whether this long-term functional re-organization of FST circuits is responsible for the clinical efficacy of psilocybin across neuropsychiatric conditions

## Materials and Methods

### Sample and study design

We leveraged longitudinal neuroimaging data and questionnaires acquired as part of a fixed-order, within-subjects design to investigate the effects of psilocybin in 25 healthy human adults with no prior psychedelic experience (***Table S1***). All participants received: (*i*) a control dose of 1 mg psilocybin on the first dosing day, considered to be a subthreshold dose unable to induce a psychedelic experience; and (*ii*) a fully active dose of 25 mg psilocybin four weeks later, considered to be a high dose inducing profound psychedelic effects (**Figure 1A**). Mental well-being was measured using the 14-item Warwick-Edinburgh Mental Wellbeing Scale (WEMWBS, population mean [range] = 51 [14-70])(50). Participants’ WEMWBS was assessed at baseline and four weeks after each dosing visit. For further information see ***Supplementary Methods*** and previous work (17).

### fMRI acquisition, preprocessing, and empirical analysis

Imaging was performed on a 3T Siemens Tim Trio using a 12-channel head coil at Imperial College London at baseline and four weeks after each dosing session. Preprocessing was conducted using standard procedures (51). Further methodological details can be found in ***Supplementary Methods*** and in previous work (17). The Automated Anatomical Labelling 90 (AAL-90) atlas (52) was then used to derive regional estimates of BOLD activity for 28 FST regions-of-interest (**Figure 1B**). The AAL was used for the following reasons: (*i*) the AAL yields excellent results in the whole-brain modelling literature (29, 30, 53); (*ii*) the relative low number of parcels in the AAL is highly suitable for our very extensive computational demands; (*iii*) the AAL atlas provides clear regional parcellations for the frontal cortex, thalamus, and basal ganglia. These standardized anatomical labels were used to select our set of FST regions-of-interest for the extraction of time series data and subsequent analyses. Intrinsic functional connectivity matrices were derived for each participant by correlating regional FST activity time series using Pearson’s correlation (***Figure S4***). DFH was estimated at the whole-brain level and for FST regions only by computing the cosine similarity in activity levels across every timepoint (***Figure S1 and S2***) (26), hence providing a measure of activity homogeneity across the duration of the entire fMRI scan. Higher values of DFH reflect higher homogeneity of brain activity in time. Mean DFH was derived by averaging the lower triangular elements of the matrix.

### Structural connectivity

Our analyses leveraged a weighted structural connectivity matrix validated and used in previous studies (24–26), derived from diffusion tensor imaging data acquired in 16 healthy young adults (5 females, mean ± standard deviation age: 24.7 ± 2.5 years). Methodological details can be found in ***Supplementary Methods*** and in previous work (24–26).

### Effective connectivity

To derive effective connectivity, we capitalized on a recently developed approach that computes the time-shifted correlations between: (*i*) forward time series of two brain regions and (*ii*) reversed time series of two brain regions (33–35). This framework was integrated within two separate whole-brain models, one for post-1 mg and another one for post-25 mg psilocybin data. In both models, we optimized effective connectivity, reflecting local constrains of structural connectivity on information flow between pairs of regions (33–35). Effective connectivity estimates were derived by running 30 separate simulations each with 1,500 iterations. The entry point for these analyses is the structural connectivity matrix obtained with probabilistic tractography from diffusion tensor imaging data in healthy volunteers and the following procedure is used to update effective connectivity:

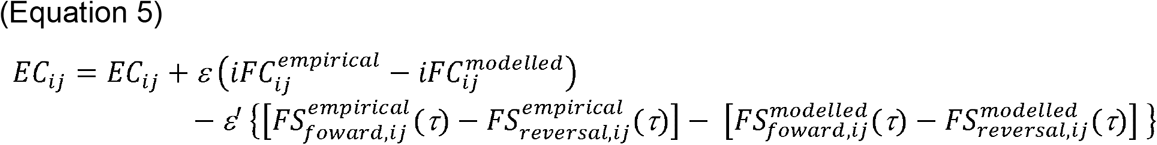

Where *EC*_*ij*_denotes the effective connectivity between pairs of regions, *iFc*_*ij*_is based on the non-shifted intrinsic functional connectivity matrices defined through the mutual information measure obtained by:

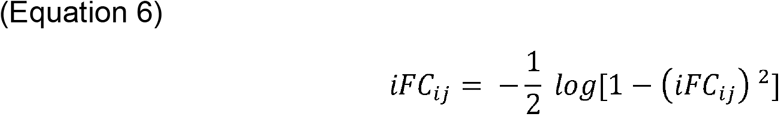

We used ε = 0.0005 and ε′ = 0.0001 and continue until the algorithm converges. *Fs*_*ij*_ reflect the time-shifted forward, *x*_*i*_ (*t*), and reversed, 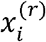, functional connectivity matrices derived with a time lag *τ* of 2 TRs (33–35):

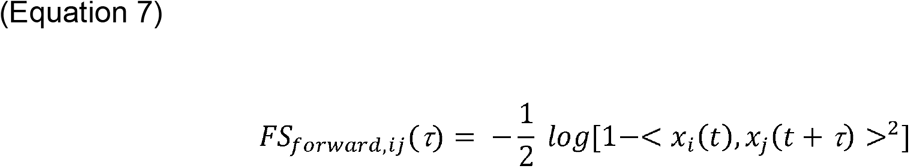

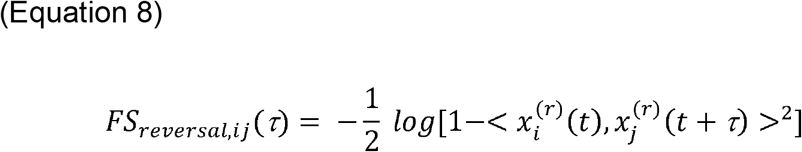

### 5-HT2A receptor and D2 receptor density maps

Publicly available PET density maps of the serotonin 5-HT2A receptor and of the dopaminergic D2 receptor were derived from a previously published study (38). Briefly, for each radiotracer, group-averaged maps were derived by only using the scans of healthy participants. Normalized PET images were parcellated to the AAL atlas and z-scored receptor density maps were derived for the 28 FST regions analyzed in this study. Detailed information on each study, the associated receptor/transporter, tracer, number of healthy participants, age, and reference with full methodological details have been previously published (38).

### Statistical analyses

MATLAB R2021a (https://www.mathworks.com/products/matlab.html) was used for statistical and computational analyses. For further details see ***Supplementary Materials***.

## Acknowledgements

We thank the volunteers of this study for their invaluable contribution to research.

## Data availability

Data is available from R.L.C.H. or L.R. after reasonable request. Scripts used to generate the data are publicly available on GitHub (https://github.com/lollopasquini).

