## Supplement for "Long-term effects of psilocybin on dynamic and effectivity connectivity of fronto-striatal-thalamic circuits"

### Content:

### Supplementary Methods

### Supplementary Figures: 5

### Supplementary Table: 1

### Supplementary References

### Supplementary Methods:

*Study sample.* This study was approved by the London-Surrey Research Ethics Committee and by the National Institute for Health Research/Wellcome Trust Imperial Clinical Research Facility. This research was carried out in accordance with Good Clinical Practice guidelines. All participants provided written informed consent. Detailed descriptions of the study sample, recruitment strategy, and of the screening procedures have been published in previous work (1). Briefly, 28 participants were initially recruited via an online advertisement on the website of the Centre for Psychedelic Research at Imperial College London. After initiating contact and following a brief telephone screen, participants were invited to a thorough in-person mental and physical examination at the National Institute for Health Research/Wellcome Trust Imperial Clinical Research Facility (ICRF) to determine eligibility. Physical assessments included: (*i*) an alcohol breathalyzer test, (*ii*) a urine screen for drugs of abuse and pregnancy (when applicable), (*iii*) an electrocardiogram, (*iv*) routine blood tests, and (*v*) blood pressure, heart rate, height, and weight recordings. Mental health assessments included (*i*) a neurological examination and (*ii*) a standard psychiatric interview, the Mini-International Neuropsychiatric Interview version 5 (2). To be included, participants had to be between 18-85 years of age, be mentally and physically healthy, speak English, and have no prior experiences with psychedelic drugs. Important exclusion criteria were a current or previous diagnosis of a psychiatric disorder, presence of a family members diagnosed with a psychotic disorder, a positive pregnancy test, and MRI contra-indications. From this initial sample, three participants had to be subsequently excluded due to high levels of head movement in the scanner and sub-optimal quality of neuroimaging data (see further details below). This procedure resulted in 25 participants (11 females, mean ± standard deviation age: 40.8 ± 8.5 years) included in this study (Supplementary Table S1).

*Study design.* We used a controlled, fixed-order, within-subjects design to investigate the effects of psilocybin in healthy human adults with no prior psychedelic experience. All participants received two oral doses of psilocybin, four-weeks apart: (i) a control dose of 1mg psilocybin on the first dosing day, considered to be a subthreshold dose that is unable to occasion a psychedelic experience; and (ii) a fully active dose of 25 mg psilocybin, considered to be a high dose and capable of inducing profound psychedelic effects, four-weeks later (Figure 1A). Bottled and encapsulated psilocybin was provided by COMPASS Pathways and securely stored at Imperial College London in light-protected and temperature-controlled conditions. This fixed order design was necessary given the hypothesized carry-over effects of 25 mg psilocybin. To uphold blinding and control for expectancy effects, participants were informed that they would receive psilocybin on both sessions of a variable dose up to 25 mg. No further information regarding dosage was provided.

*Neuroimaging data acquisition.* Imaging was performed on a 3T Siemens Tim Trio using a 12-channel head coil at Imperial College London. Whole-head anatomical images were acquired using the Alzheimer’s Disease Neuroimaging Initiative, Grand Opportunity (ADNI-GO)(3) recommended Magnetization Prepared Rapid Gradient Echo (MPRAGE) parameters – 1 mm isotropic voxels, TR = 2300 ms, TE = 2.98 ms, 160 sagittal slices, 256 × 256 in-plane FOV, flip angle = 9°, bandwidth = 240 Hz/pixel, parallel imaging (PI) factor = 2, Inversion time = 900 ms. For resting-state fMRI, T2*-weighted echo-planar images (EPI) were acquired with interleaved slice acquisitions using 3 mm isotropic voxels, TR = 1250 ms, TE = 30 ms, 44 axial slices, 192 mm in-plane FOV, flip angle = 80°, bandwidth = 2232 Hz/pixel, GRAPPA acceleration = 2, number of volumes = 384. Resting-state scans lasted 8 minutes and were completed with eyes-closed.

*Neuroimaging data preprocessing.* fMRI data was preprocessed as described in previously published work (1). FMRIB Software Library (FSL)6, Analysis of Functional NeuroImages (AFNI)7, Freesurfer 8, and Advanced Normalization Tools (ANTS)9 were used to analyze the resting-state data. 10. The following preprocessing stages were performed: (*i*) removal of the first three volumes; (*ii*) de-spiking (3dDespike, AFNI); (*iii*) motion correction (3dvolreg, AFNI) by registering each volume to the volume most similar, in the least squares sense, to all others (in-house code); (*iv*) brain extraction (BET, FSL); (*v*) rigid body registration to anatomical scans (FSL, BBR); (*vi*) non-linear registration to 2mm MNI brain (Symmetric Normalization (SyN), ANTS); (*vii*) scrubbing 11 using an FD threshold of 0.4 mm. Scrubbed volumes were replaced with the mean of the surrounding volumes. Additional preprocessing steps included: (*viii*) spatial smoothing (FWHM) of 6 mm (3dBlurInMask, AFNI); (*ix*) band-pass filtering between 0.01 to 0.08 Hz (3dFourier, AFNI); (*x*) linear and quadratic de-trending (3dDetrend, AFNI); (*xi*) regressing out 9 nuisance regressors (the same bandpass filter was applied on the nuisance regressors): out of these, 6 were motion-related (3 translations, 3 rotations) and 3 were anatomically-related (not smoothed). Specifically, the anatomical nuisance regressors were: (a) ventricles (Freesurfer, eroded in 2mm space), (b) draining veins (DV) (FSL’s cerebrospinal fluid minus Freesurfer’s Ventricles, eroded in 1mm space) and (c) local white matter (WM) (FSL’s WM minus Freesurfer’s subcortical grey matter (GM) structures, eroded in 2mm space). Regarding local WM regression 12, AFNI’s 3dLocalstat was used to calculate the mean local WM time-series for each voxel, using a 25mm radius sphere centered on each voxel. Motion was measured using frame-wise head displacement. Subjects were included only if their mean frame-wise head displacement in the scanner (38,39) was below 0.8 mm, resulting in the final sample of 25 subjects analyzed in this study (40). Mean framewise head displacement did not significantly differ between fMRI scans (**Table S1**) (4, 5).The Automated Anatomical Labelling 90 (AAL-90) atlas (6) was then used to derive regional estimates of BOLD activity for 28 FST regions-of-interest (**Figure 1B**). The AAL was used for the following reasons: (*i*) AAL yields excellent results in the whole-brain literature in general (7–9); (*ii*) the relative low number of parcels in the AAL is highly suitable for our very extensive computational demands; (*iii*) the AAL atlas provides clear regional parcellations for the frontal cortex, thalamus, and basal ganglia. These standardized anatomical labels were used to select our set of FST regions-of-interest for the extraction of time series data and subsequent analyses. The derived FST activity time series were subsequently bandpass filtered in the 0.04 to 0.08 Hz frequency range. This frequency band has been shown to be less susceptible to artefactual sources such as low-frequency drift, cardiac and respiratory variations, and to preferentially map on gray matter instead of white matter and cerebrospinal fluid (10, 11). Further, removing the lower-frequency range has been shown to improve the correspondence between neural activity and hemodynamic signals based on simultaneous fMRI/electrophysiological recordings in awake rats (12). Finally, this frequency range has been widely used in previous studies using whole-brain models, yielding positive outcomes (13–16).

*Structural connectivity*. A whole-brain and FST structural connectivity matrices were derived for modelling purposes by leveraging a structural connectivity matrix validated and used in previous studies (9), which leveraged diffusion tensor imaging data acquired in 16 healthy young adults (5 females, mean ± SD age: 24.7 ± 2.5 years). The FSL diffusion toolbox was used to preprocess this data as described in previous work (9) and the local distribution of fiber direction at each voxel was estimated. For each brain region, the connectivity probability to each of the other 89 regions within the AAL was calculated. As directionality of connections cannot be determined based on diffusion MRI, the unidirectional connectivity probability *P_ij_* between regions *i* and *j* was defined by averaging these two connectivity probabilities. Connectivity probability values were then normalized by the number of voxels in each AAL region. This whole-brain matrix was used for modelling effective connectivity. Values of the 28 FST regions were extracted resulting in a symmetric weighted matrix of FST structural connectivity which was used in the partial-brain models to estimate the *G* coupling parameter. For diffusion tensor imaging (DTI) the following parameters were used: TR = 9000 ms, TE = 84 ms, flip angle = 90°, reconstructed matrix size of 106×106, voxel size of 1.98×1.98 mm, slice thickness = 2 mm and bandwidth = 1745 Hz/Px. Data were recorded using 62 optimal nonlinear diffusion gradient directions at b = 1500 s/mm². For every 10 diffusion-weighted images, one non-diffusion-weighted image (b=0) was acquired. The first diffusion MRI image was collected with an anterior to posterior phase encoding direction, while the second image was acquired in the opposite direction (8).

*Partial-brain model.* Whole-brain Hopf models are used to model the dependency of emerging functional activity dynamics from the underlying structural brain connectivity (17–19). Here, we used the supercritical Hop-bifurcation model for a single uncoupled region of interest j to derive a partial-brain model, which, instead of focusing on the whole brain, consists of coupled dynamical units of FST regions only. The dynamics of a brain region *n* is given by the following set of coupled dynamical equations:

(Equation 1)

$$\frac{{dx}_{j}}{dt}=\left[ a_{j}-x_{j}^{2}-y_{j}^{2} \right]x_{j}- \omega_{j}y_{j}+{\beta\eta}_{j}(t)$$

(Equation 2)

$$\frac{{dy}_{j}}{dt}=\left[ a_{j}-x_{j}^{2}-y_{j}^{2} \right]y_{j}+ \omega_{j}x_{j}+{\beta\eta}_{j}(t)$$

where ${\beta\eta}_{j}(t)$ is additive Gaussian noise with standard deviation $\beta=0.01$. This normal form has a supercritical bifurcation $a_{j}$= 0, so that if $a_{n}$> 0, the system engages in a stable limit cycle with frequency $f_{j}= \omega_{j}/2\pi$. On the other hand, when $a_{j}$< 0, the local dynamics are in a stable fixed point representing a low activity noisy state. Within this model, the intrinsic frequency is estimated from the empirical data as the peak of the power spectrum between 0.04 and 0.08 Hz (***Figure S5***).

To model the partial-brain dynamics, we added an additive coupling term representing the input received in node *j* from every other node *i*, which is weighted by the corresponding structural connectivity represented by $C_{ij}$. The partial-brain dynamics were defined by the following set of coupled equations:

(Equation 3)

$$\frac{{dx}_{j}}{dt}=\left[ a_{j}-x_{j}^{2}-y_{j}^{2} \right]x_{j}- \omega_{j}y_{j}+G\sum_{i=1}^{N} C_{ij}\left( x_{i}- x_{j} \right)+ {\beta\eta}_{j}(t)$$

(Equation 4)

$$\frac{{dy}_{j}}{dt}=\left[ a_{j}-x_{j}^{2}-y_{j}^{2} \right]y_{j}+ \omega_{j}x_{j}+G\sum_{i=1}^{N} C_{ij}\left( y_{i}- y_{j} \right)+ {\beta\eta}_{j}(t)$$

where the noise standard deviation $\beta$= 0.01. The local bifurcation parameters, $a_{j}$= -0.02, are at the brink of the local bifurcations which is where the best fitting was demonstrated to be achieved in previous work (17–19). The variable $x_{j}$ emulates the BOLD signal of each FST region *j*. The term *G* denotes the global coupling weight, scaling equally the total input received in each brain area. Two partial-brain models were separately estimated for the post-1 mg and post-25 mg conditions for varying values of the global coupling parameter *G* by fitting simulated and empirical DFH at each step. For both models, we run 30 separate simulations each with 1,500 iterations.

*Statistical analyses.* Repeated measures ANOVA and associated paired t-tests were used to compare mean frame-wise head displacement in the scanner and changes in WEMWBS across scanning and dosing sessions. Paired t-tests were used to assess changes in mean DFH cosine similarity of FST regions across the three time points. Kolmogorov-Smirnov statistics were used to estimate partial-model fits by comparing the distributions of modelled DFH cosine similarity to empirical DFH cosine similarity. The correspondence of modelled and empirical intrinsic functional connectivity matrices was estimated through Pearson’s correlation. Wilcoxon rank sum tests were used to compare fit estimates from the partial-brain model applied to data from the second and third scan. Paired t-tests were used to assess statistical changes in effective connectivity taking place between the second and third scan. Paired t-tests were also used statistically compare absolute values of mean asymmetry in effective connectivity between the second and third scan. Spearman correlation was used to associate changes in WEMWBS scores between the third and second scan to change in mean DFH cosine similarity in the same time interval. This analysis was corrected for baseline well-being and head movement in the scanner by regressing out both the influence of baseline WEMWBS scores and change in mean head frame-wise displacement between both scan sessions. Spearman correlation coefficients were further used to spatially associate regional estimates of the 5-HT2A receptor and of the D2 receptor with nodal estimates of changes in incoming and outcoming effective connectivity following 25 mg psilocybin. Here, separate analyses were performed for cortical and subcortical FST regions due to the marked differences in receptor and transporter densities when comparing subcortical and cortical areas. Control analyses were performed by associating whole-brain changes in effective connectivity to whole-brain receptor maps. Fisher’s r to z transformation was used to significantly compare the strength in correlation between subcortical and cortical areas. MATLAB R2021a (<https://www.mathworks.com/products/matlab.html>) was used for statistical and computational analyses.

### Supplementary Figures, Legends, and Tables:

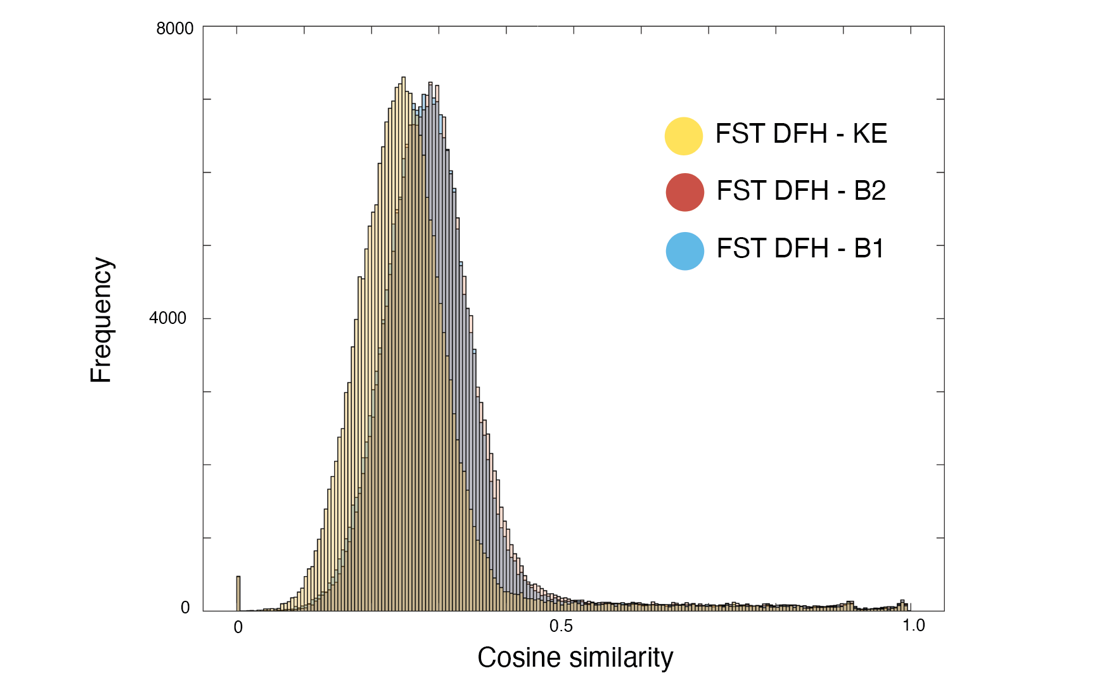

**Supplementary** **Figure S1. FST dynamic functional homogeneity distribution.** Group-level FST dynamic functional homogeneity (DFH) distributions for B1 (in blue), B2 (in red), and KE (in yellow) generated by using FST regions only. Note the large overlap between the DFH distributions of B1 and B2, while the DFH distributions of KE are shifted towards the left, indicating lower DFH following 25 mg psilocybin.

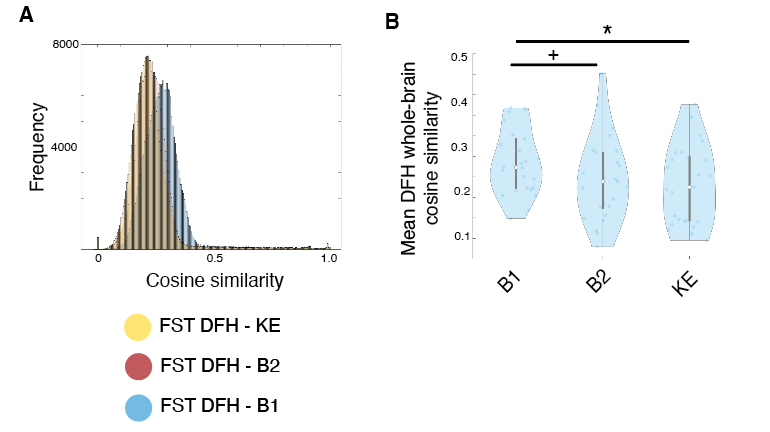

**Supplementary** **Figure S2. Whole-brain dynamic functional homogeneity.** Group-level whole-brain dynamic functional homogeneity (DFH) distributions for B1 (in blue), B2 (in red), and KE (in yellow) generated by using all AAL90 regions. Note the large overlap between the DFH distributions of KE and B2, while the DFH distributions of B1 are shifted towards the right, indicating higher whole-brain DFH at baseline. (**B**) Group differences in mean whole-brain DFC between the three scanning time-points. **p* < 0.05; +*p* < 0.01

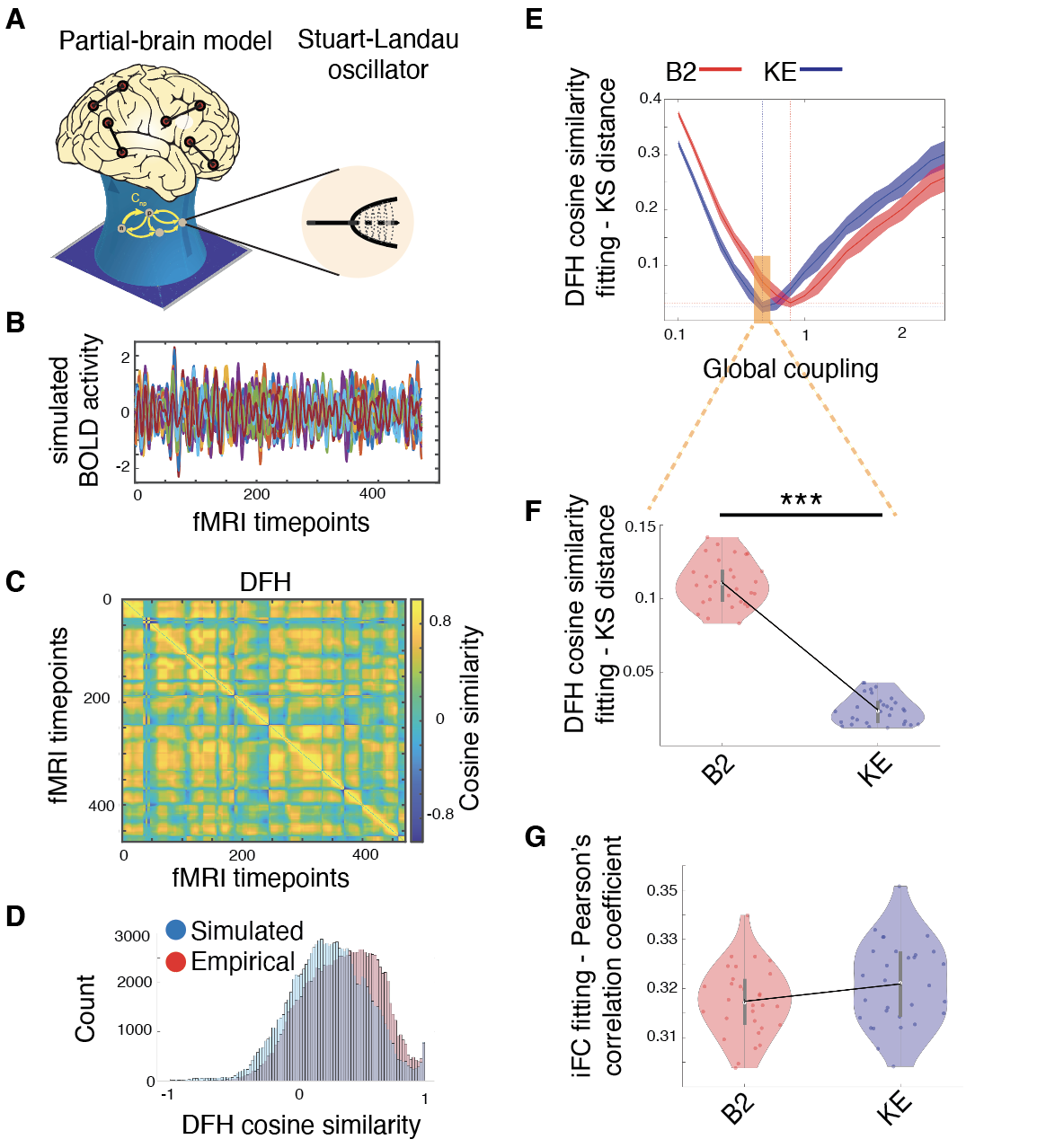

**Supplementary** **Figure S3. Reduced global structure-function coupling after 25 mg psilocybin. (A)** We derived a partial-brain model of FST dynamics by using a Stuart-Landau oscillatory system. This system describes a set of differential equations defined by two major parameters: (*i*) a bifurcation parameter *a* describing the propensity of a system to display noisy or oscillatory dynamics, which is kept constant ($a_{j}$= -0.02); and (*ii*) a global coupling parameter *G* weighting the influence of underlying structural connectivity on functional dynamics. Low or high global coupling parameters reflect whether functional networks are weekly or strongly constrained by the underlying structural connections. **(B)** The partial-brain model was used to iteratively generate simulated BOLD activity time series that best reproduced the statistical properties of empirical FST DFH acquired at B2. **(C)** Example of a simulated DFH cosine similarity matrix derived from modelled FST activity time series. **(D)** Distributions of empirical (in red) and simulated (in blue) DFH values were compared using Kolmogorov-Smirnov (KS) distance to select the partial-brain model providing the best fit. **(E)** Model fitting to empirical DFH as a function of the global coupling parameters plotted separately for B2 (in red) and KE (in blue). Model fitting values were derived from 30 separate converging simulations. Solid line and shade reflect the average and standard deviation, respectively. **(F)** Significantly improved model fitting for KE versus B2 when using the optimal global coupling parameter of KE to fit DFH. **(G)** Model fit for intrinsic functional connectivity (iFC), as indexed through a Pearson’s correlation coefficient, did not significantly differ between KE and B2, suggesting that fitting DFH is more sensitive to changes in structure-function coupling than static iFC. Panel A adapted with permission from (17). BOLD = blood-oxygen-level-dependent. ****p* < 0.0005.

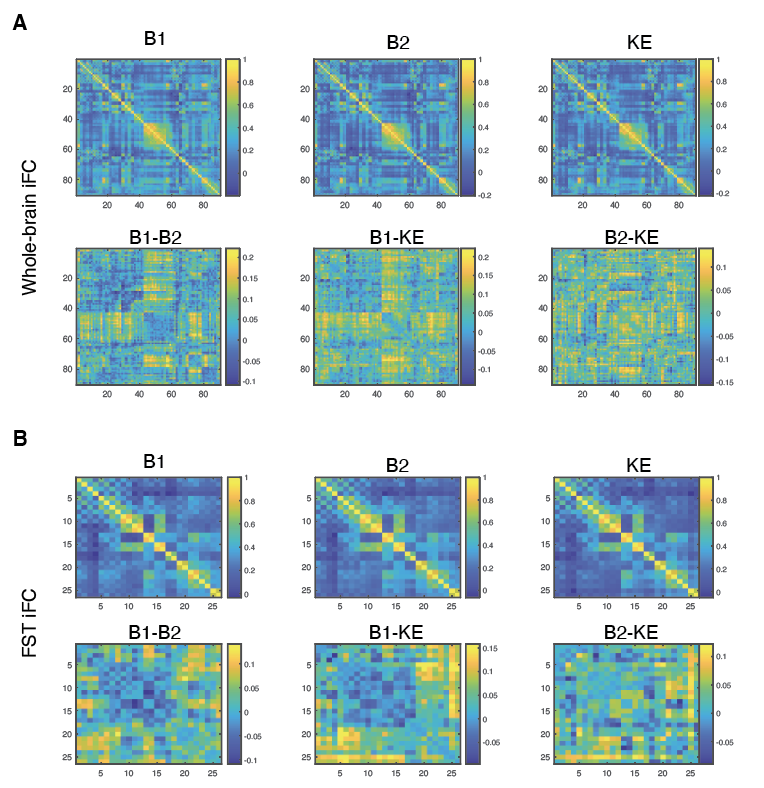

**Supplementary** **Figure S4. Group-mean intrinsic functional connectivity matrices.** Group-mean intrinsic functional connectivity matrices for B1, B2, and KE and associated mean-group differences once at the whole-brain level (**A**) and for FST regions only (**B**). Axes in the heatmaps represent brain regions.

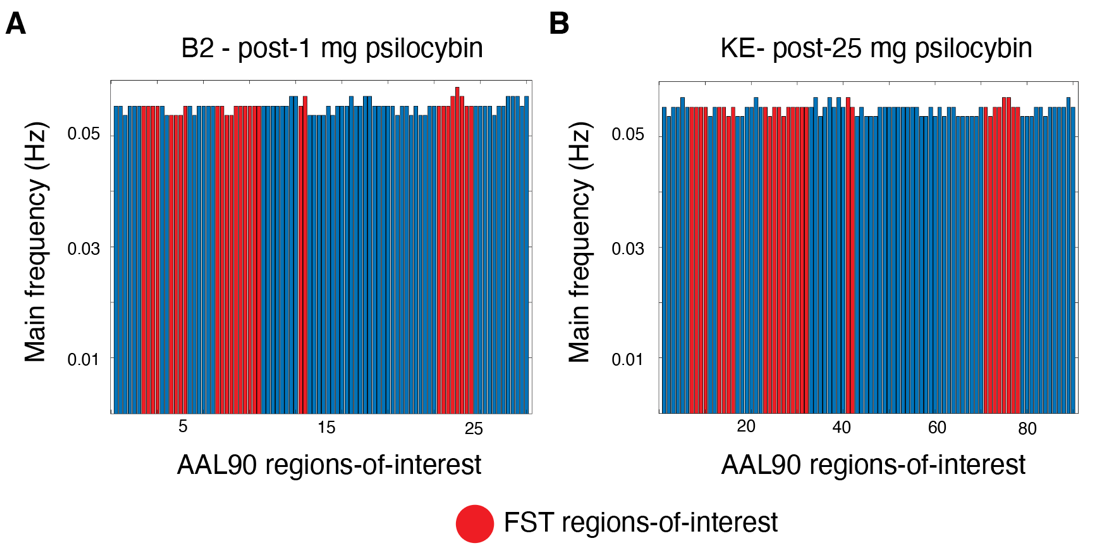

**Supplementary** **Figure S5. Main intrinsic frequency.** Main intrinsic frequency (in Hz) for AAL-90 regions-of-interests estimated for the post-1 mg psilocybin condition (B2, **A**) and for the post-25 mg psilocybin condition (KE, **B**). FST regions-of-interest are shown in red bars.

|  | **N=25** | ***F*** | ***p*** |
| --- | --- | --- | --- |
| *Baseline assessment* | | | |
| Age in years, mean (SD; range) | 40.8 (8.5; 28-59) |  | |
| Gender (female/male) | 11/14 |  |  |
| Education (secondary/university) | 11/14 |  |  |
| Race (Black/White/Undisclosed) | 1/21/3 |  |  |
| Nationality (British/Other) | 19/6 |  |  |
| *Longitudinal assessment* | | | |
| WEMWBS – B1 (n=24), mean (SD) | 51.7 (6.3) | 6.04 | <0.005 |
| WEMWBS – B2 (n=24), mean (SD) | 49.3 (6.5) |  |  |
| WEMWBS – KE (n=25), mean (SD) | 54.0 (6.7) |  |  |
| Mean FD in mm – B1, mean (SD) | 0.14 (0.11) | 1.22 | *p*=0.30 |
| Mean FD in mm – B2, mean (SD) | 0.15 (0.13) |  |  |
| Mean FD in mm – KE, mean (SD) | 0.16 (0.12) |  |  |

**Supplementary Table S1.** Demographic characteristics of the analyzed sample. FD = mean frame-wise head displacement; WEMWBS = Warwick-Edinburgh Mental Wellbeing Scale
